# What Primates Know About Other Minds and When They Use It: A Computational Approach to Comparative Theory of Mind

**DOI:** 10.1101/2023.08.02.551487

**Authors:** Marlene D. Berke, Daniel J. Horschler, Amanda Royka, Laurie R. Santos, Julian Jara-Ettinger

## Abstract

Can non-human primates (NHPs) represent other minds? Answering this question is difficult because primates can fail tasks due to a lack of motivation or succeed through simpler strategies. Here we address these challenges through a computational theory-testing framework for NHP Theory of Mind. In this framework, each theory combines a proposed social representation with a parameter for how often it is used. This allow us to move beyond dichotomous positions about Theory of Mind’s presence or absence and instead analyze graded patterns of behavior as a combination of cognitive representations and their use. We apply this approach to one of the most foundational and well-studied aspects of Theory of Mind: the relation between seeing and knowing. Our results show that only theories in which NHPs have some representation of other minds can capture the qualitative pattern of successes and failures across five classic perspective-taking paradigms. However, these theories vary in their reliance on their representations, each showing significantly lower reliance than a human baseline. These results suggest that human and NHP social cognition differ in terms of reliance and possibly also in terms of representational complexity.

---

Like humans, non-human primates (NHPs) have rich social lives: they live in complex social groups, can act altruistically, show empathy for group members, and even develop lifelong social relations—from dominance hierarchies to close friendships (Warneken et al., 2007; Lewis et al., 2023; Muller & Mitani, 2005; Cheney & Seyfarth, 2008). At the same time, their social behavior is undeniably simpler than that of humans (Hall & Brosnan, 2017; Hoppitt et al., 2008; Musgrave et al., 2016; Tomasello et al., 2005): NHPs do not intentionally deceive one another, they only engage in limited forms of pedagogy, and they do not develop cultures and societies of human-level complexity. What cognitive differences might explain this gap?

Part of the answer is thought to lie in *Theory of Mind*—the ability to interpret and predict others’ behavior by representing unobservable mental states like beliefs and desires. Theory of Mind supports a broad range of human capacities, including language use, cooperation, and moral reasoning (Gweon, 2021; Rubio-Fernandez et al., 2024; Young et al., 2007). Its foundational components emerge early in infancy (Csibra et al., 2003; Liu et al., 2017; Gergely & Csibra, 2003; Repacholi & Gopnik, 1997), and many important mental-state inferences become automatic in adults (Kampis & Southgate, 2020; Rubio-Fernández et al., 2019). As such, differences between human and NHP Theory of Mind (hereafter NHP ToM) might explain the gaps in the relative complexity of social life.

This hypothesis has led to extensive research on comparative Theory of Mind (see Rosati et al., 2010; Krupenye & Call, 2019; Call & Tomasello, 2008, for reviews), but there is little consensus on what exactly NHPs understand about others’ mental states (compare, e.g., Martin & Santos, 2016; Krupenye et al., 2016). This stems from two fundamental challenges in comparative research.

The first challenge is practical. When NHPs fail in a ToM task, is it because they lack this capacity, or because they did not *rely* on it? That is, NHPs might be able to represent their conspecifics’ minds, but fail to do so in a particular task. This could happen for a variety of reasons, such as the cognitive load that a task imposes on them (e.g., via time pressure or task complexity) or their motivation to understand their conspecifics’ behavior in that particular situation. For example, some research has argued that NHPs have a ToM, but they don’t always rely on it (Hare & Tomasello, 2004; Kano et al., 2020). Eliciting it may require adversarial tasks or situations with “social drama.” This idea that NHPs do not consistently rely on their social representations helps explain why only some tasks elicit ToM-like behavior and why these tasks sometimes produce weaker effect sizes than one would expect in humans.

The second challenge is theoretical. Success on ToM tasks can often be explained by non-mentalistic, rule-based accounts. This is a common challenge in comparative research—for nearly every study on Theory of Mind, there exist competing mentalistic and non-mentalistic accounts that explain the qualitative data equally well (Penn & Povinelli, 2007b; Andrews, 2012, 2018; Heyes, 1998).

Our paper seeks to introduce a computational approach that can help differentiate between competing accounts of NHP social cognition. The high-level goal is to implement a family of computational models, each positing a different representation that NHPs might use in social tasks. Each account, by virtue of its implementation as a computational model, not only predicts qualitative results (such as whether a NHP might succeed or fail in a task) but also quantitative effect sizes that can be directly compared to NHP experimental data.

Our computational framework is centered around these challenges in interpreting NHP behavior in social tasks. Failure, on the one hand, might mean that NHPs were not relying on their available social representations. Success, on the other hand, might be explained through non-mentalistic rule-based accounts. These challenges are often treated as independent from each other, but we propose that they are intrinsically connected. Intuitively, the complexity of the social representation that we posit should be related to the frequency with which NHPs rely on it. For instance, if NHPs rely on a trivial behavioral rule, we might expect them to show near ceiling performance on the task, whereas a complex mentalistic representation, in some cases, can create more uncertainty for the NHP, predicting a weaker effect size.

Given a theoretical account of NHP social reasoning, our approach is to implement it as a computational model that predicts effect sizes across a set of experiments. Usually, the gap between predicted and observed effect sizes determines a theory’s plausibility. But this evaluation assumes that all subjects always rely on the posited representation. To avoid this, we introduce a parameter for reliance that helps explain observed discrepancies. This allows us to use the difference between predicted and observed effect sizes to estimate how often NHPs are relying on the theory’s posited social representations if the account were true. This reveals the hidden link between different proposals about NHP social representations and the frequency with which subjects must use them for the theories to hold.

For example, if a theory predicts a strong effect but the experiment reveals a weak one, then the theory can only be true if NHPs use the representation some of the time. This is not necessarily a flaw, and might even be a desirable theoretical property of an account, since NHPs might not always rely on complex or costly representations. Thus, our computational framework serves as a tool to clarify these implicit commitments that different theories entail. Critically, not all discrepancies can be explained by reliance. While reliance can account for effect sizes that are weaker than predicted, it cannot explain explain cases where the observed effect is stronger than what a theory predicts. If a theory predicts a weak effect size, but NHPs show near-ceiling performance, this suggest that NHPs are using a different representation that helps them to perform so well in the task. This means that our approach is also able to discard theories when they under-predict how NHPs behave.

Using this approach, our goal is to propose a methodology for formalizing competing theories of NHP social representations and estimate expected effect sizes in a way that can be directly compared to published NHP studies. We then show how our approach allows us to extract the hidden assumptions about reliance that different theories make by examining the difference between predicted and observed effect sizes.

Our approach differs from previous work applying computational modeling to NHP social cognition (e.g., communication and coordination; Bohn et al. 2022; Nong 2023). While important, past work has primarily focused on components of NHP behavior where computational models of human behavior can be applied to NHPs—i.e., cases where the conclusion is that NHPs think and behave like humans. Our approach is complementary, focusing instead on how to formalize multiple competing theories of mental representation in NHPs and evaluate their explanatory power.

Because Theory of Mind is a complex cognitive system with many sub-components (including mechanisms for inference, prediction, explanation, and even planning over other minds; Ho et al. 2022; Wang et al. 2025), here we focused on just one component of Theory of Mind that has been well-researched in comparative cognition: the ability to determine what others see and know based on their visual perspective (see Phillips et al., 2020, for review). The capacity to represent the link between seeing and knowing is one of the most basic components of ToM, emerging early in human infancy (Luo & Johnson, 2009), and the question of whether these capacities are shared with our NHP relatives has been the subject of much debate (Call & Tomasello, 2008; Halina, 2017; Penn & Povinelli, 2007a; Povienlli & Vonk, 2003; Rosati et al., 2010; Lurz et al., 2014).

To illustrate this capacity, consider Fig. 1a, which shows a subordinate chimpanzee (the subject; black) and a dominant conspecific (the competitor; gray) on opposite sides of a room with two apples. One apple is in the center of the room and visible to both chimpanzees, while the second apple is behind an opaque barrier and visible only to the subject. If the subject understands that the competitor only knows about the apple in the center of the room, they can use this to strategically decide which apple to pursue. Tasksb like this were used in now-classical work to show that chimpanzees (*Pan troglodytes*) will preferentially approach food rewards that are hidden from conspecifics, providing some of the first evidence for visual perspective-taking in NHPs (Hare et al., 2000). This work has since been extended to show more complex forms of perspective-taking, including evidence that NHPs know what others can hear (Santos et al., 2006; Melis et al., 2006).

**Figure 1.**
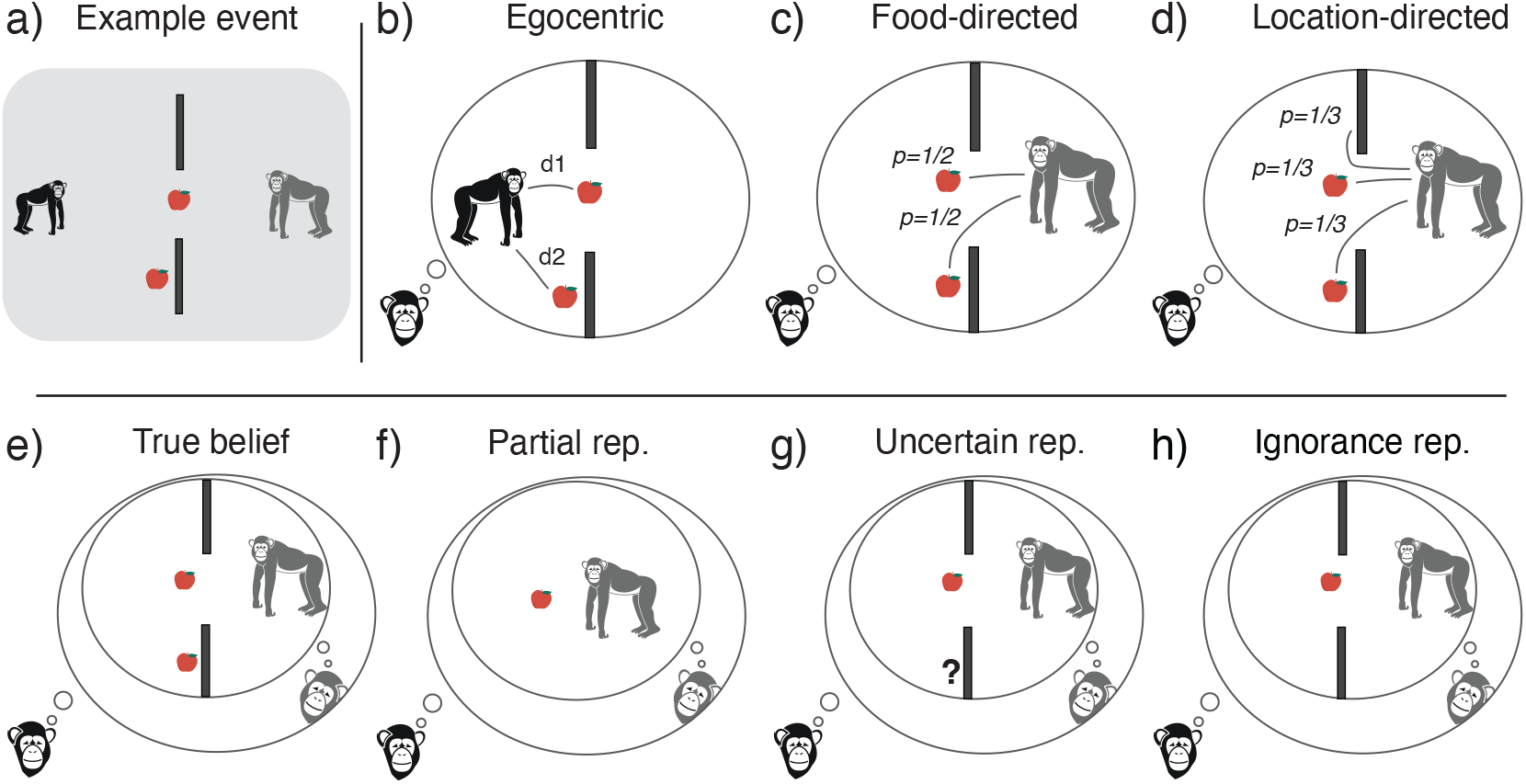
a) A hypothetical competitive task used to test the relation between seeing and knowing. Here, the subject (small chimpanzee, black) is in front of a dominant conspecific (big chimpanzee, gray) in a room with two apples and two opaque barriers, such that both chimpanzees see the apple between them, but only the subject sees the apple in front of the barrier. b-h) Visual illustration of our seven computational models, ordered by complexity. The models fall into three broad families: non-mentalistic (b-d), semi-mentalistic (e-g), and full ToM (h). Throughout, p refers to probabilities, and d refers to distances.

To explore possible representations, we developed seven computational models of varying cognitive complexity (shown in Fig. 1b-h and Table 2; explained in detail in the following section). Each model posits a different cognitive representation and is instantiated as a simple system that can complete basic primate experiments implemented in two-dimensional grid worlds. This allowed us to implement competing theories of NHP social cognition as computational models and compare their performance against NHP’s performance on visual perspective-taking tasks. We then included a component that captures how often NHPs rely on their representations, which we call *reliance*. Our construct of reliance is intentionally broad, designed to capture the frequency with which NHPs use the posited representation, without a commitment to the specific causes behind it. Thus, reliance can be thought of as an aggregate measure that captures several reasons why NHPs might not use their representations: for example, the task might not be insufficiently motivating or it may pose too large a cognitive load (e.g., due to task complexity or time pressure).

Our framework makes two key contributions to the study of comparative Theory of Mind. First, by developing a theory-testing framework for the visual perspective-taking literature, we offer a systematic test of different representational accounts of these capacities. Specifically, we evaluated each model on a suite of seminal visual perspective-taking tasks and compared its pattern of successes and failures to those documented in comparative studies. This approach reveals which models capture the qualitative pattern of results (i.e., when a model’s predictions of successes and failures align with empirical findings) and which levels of reliance maximize their explanatory power. Second, our framework is not limited to visual perspective-taking, and it can be applied to other debates within comparative cognitive science, both within ToM (e.g., the nature of NHP belief-like representations) and beyond (e.g., NHP intuitive physics). In this way, we hope to lay the groundwork for a broader use of computational methods in comparative cognitive science. Critically, this approach goes beyond applying computational frameworks from human cognition; instead, it is tailored to the unique challenges of comparative research. It focuses on formalizing and testing variations of theories that make the same qualitative predictions and on uncovering the implicit assumptions that each theory makes about representational reliance.

## Overview of Experimental Paradigms and Modeling

### Experimental paradigms

Our work focused on eleven experiments and controls designed to assess non-human primates’ (NHPs) understanding of how another agent’s visual perspective affects their knowledge (Table 1). These eleven experiments fall into five general paradigms (shown in Fig. 2) which follow a similar structure: A subject and a dominant competitor face one another on opposite sides of an enclosure with two food rewards that they ultimately compete for. Because dominant individuals in NHP species tend to monopolize food and respond antagonistically to challenges from lower-ranking individuals (Noë et al., 1980), these paradigms create a pressure for the subordinate subject to exploit any privileged knowledge of food locations to ensure they obtain a reward.

**Table 1.** Papers and experiments used to evaluate our computational models. Note that Hare et al. (2000) Exp 3 and 4 had multiple conditions that used more than one paradigm and are therefore repeated. All experiments tested chimpanzees (Pan troglodytes) except for Canteloup et al. (2016), which tested Tonkean macaques (Macaca tonkeana).

| Paradigm | Paper | Exp |
| --- | --- | --- |
| Center-Wall | Hare et al. (2000) | E1 |
| Open-Hidden | Bräuer et al. (2007) | E2 |
|  | Canteloup et al. (2016) | E1 |
|  | Hare et al. (2000) | E2-E4 |
| Transparent-Hidden Routes | Hare et al. (2006) | E2 |
|  | Melis et al. (2006) | E1 |
| Hidden-Hidden | Hare et al. (2000) | E3-E4 |
| Open-Transparent | Hare et al. (2000) | E5 |

**Figure 2.**
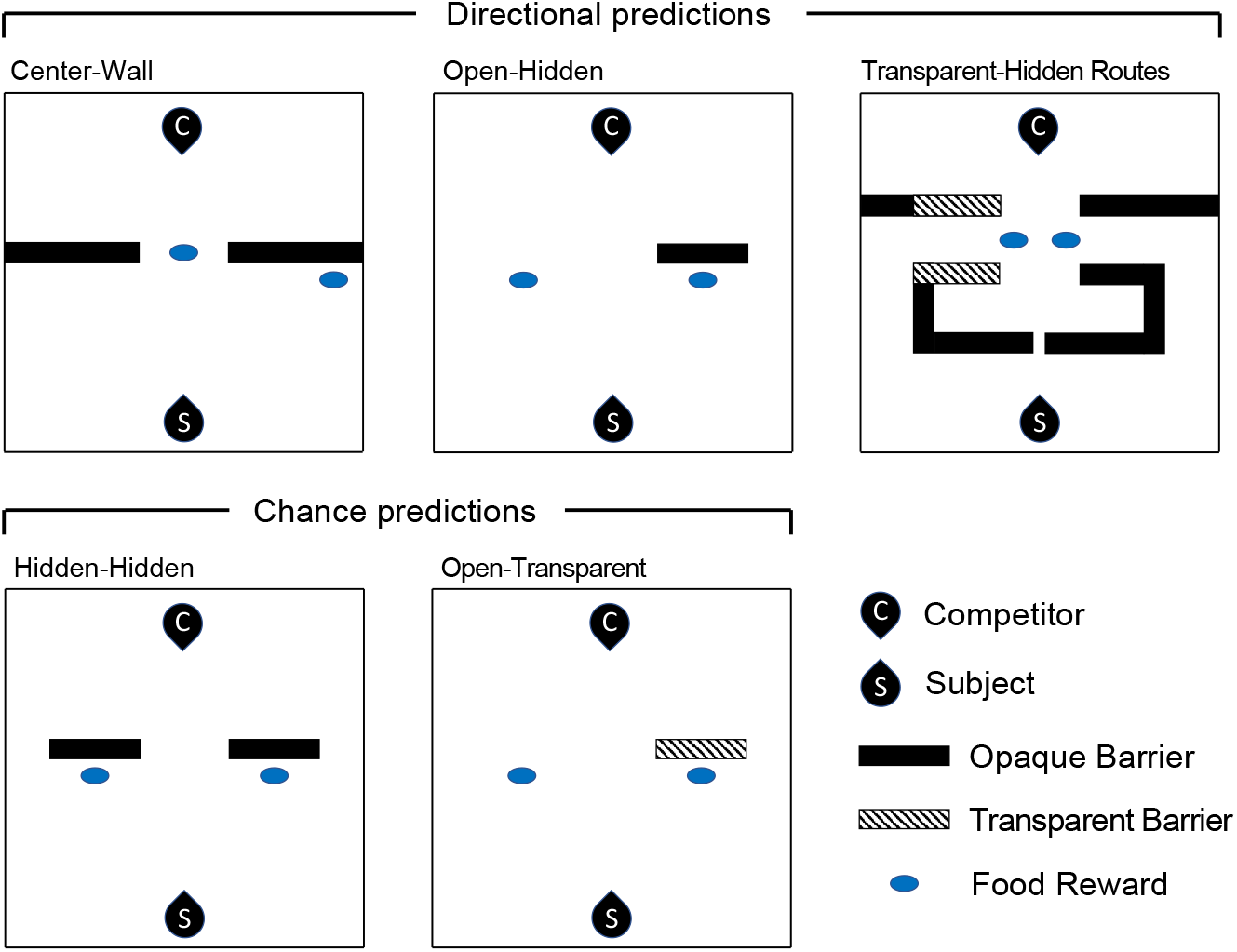
Depictions of the experimental setup for each of the five paradigms we used to compare our computational models to NHP behavior. The top row shows paradigms with directional predictions: if subordinate NHPs represent others’ knowledge, then they should preferentially go towards the reward (or take the route) that is hidden from the competitor. The bottom row shows control paradigms. When the two rewards are simultaneously known or hidden from the competitor, the subject should show no reliable preference for which one to pursue.

In the Center-Wall paradigm, the subject and the dominant conspecific face each other with a visible food reward directly between them, but the subject can also see a second food reward that is hidden from the competitor’s view by a barrier. The Open-Hidden paradigm is similar, with the difference that the two rewards are equidistant from the subject. In both of these paradigms, NHP subjects are significantly more likely to take the hidden reward. In the Transparent-Hidden Routes paradigm, the subject can reach for a food reward through a left or right path, but one of the paths is hidden from the competitor’s sight for longer than the other. Here, NHPs are more likely to reach through the hidden route than through the transparent one.

The final two paradigms are controls. In the Hidden-Hidden paradigm, both food rewards are visible to the subject but hidden from the competitor’s perspective. In the Open-Transparent paradigm, the situation is identical to the Open-Hidden paradigm except that the barrier is transparent (so the subject and competitor see both food rewards). In both of these paradigms, NHPs show no systematic preference for either food reward.

### Computational framework

To test what social representations underlie NHP behavior in these tasks, we implemented seven computational models that vary in representational complexity. These models are not meant to be faithful representations of existing theories. This is because verbal theories, while powerful, lack the details necessary to implement an exact computational model. Only the proposers of these theories can specify the exact computational implementation that would capture their nuanced positions, and we return to this point in the general discussion. Instead, the theories we present are meant to be a representative sample from the space of possible theories, varying across levels of complexity. All model and analysis code are available in our OSF repository: https://osf.io/qjn9m/?view_only=a8ede1cffb684452af9b2c54cb5ac15c.

Given a set of possible rewards that the NHP can pursue (see Fig. 2 for paradigms), each model aims to predict the probability that an NHP will choose to approach reward *R*, given by

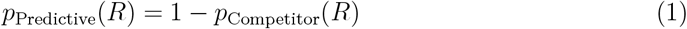

where *p*_Competitor_(*R*) is the probability that the competitor will also go for reward *R* according to the subject’s cognitive representations. This equation captures the idea that the subject wants to avoid the reward that the dominant conspecific will choose. The subject will choose reward *R* with probability 1 only when certain that the competitor will not go for it (i.e., *p*_Competitor_(*R*) = 0). Conversely, the subject will never choose reward *R* if confident that the competitor will go for it (i.e., *p*_Competitor_(*R*) = 1). This is the general formulation of all models, with each model specifying a different mechanism through which the NHP predicts the competitor’s choice *p*_Competitor_.

In some cases, NHPs might not compute or rely on *p*_Competitor_(*R*) at all. In those cases, we assume that the NHP acts egocentrically, as they would in the absence of a competitor, denoted by *p*_Egocentric_(*R*). The propensity to predict the competitor’s behavior is determined by *reliance*, which we denote as *r*. In our formulation, the NHP’s proclivity to choose reward *R* is given by

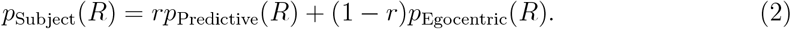

According to this equation, the NHP acts based on its prediction of the competitor’s behavior (specified in Eq. 1) with probability *r* (reliance). Otherwise (with probability 1 −*r*), the NHP acts egocentrically, without integrating predictions about the competitor.

Thus, in this formulation, reliance *r* is a free parameter that can be fit to NHP data, but its value will depend on which mechanism we posit for predicting competitor behavior. This means that, each theory of primate social representations produces a different *p*_Predictive_(*R*) estimate (by predicting different competitor behavior), and we then adjust the reliance to maximize the model’s alignment with the empirical data.

In the remainder of this section, we first discuss how we compute egocentric behavior, *p*_Egocentric_(*R*), and then go through the models, each specifying a different mechanism for predicting the competitor’s behavior *p*_Competitor_(*R*).

### Modeling egocentric behavior

To compute *p*_Egocentric_(*R*), we implemented the experimental paradigms into a Markov Decision Process (MDP; Bellman, 1957). MDPs are a general framework for modeling how agents take sequential actions in an environment to obtain rewards. Because MDPs are a popular framework in cognitive science and social cognition, here we assume some basic knowledge of this framework. A complete introduction to MDPs can be found in Sigaud & Buffet (2013), and a tutorial for MDPs applied to social reasoning can be found in Jara-Ettinger et al. (2024).

Using MDPs, we represent each experimental paradigm as a simple grid world where the NHP can sequentially move in any of the four cardinal directions (up, down, left, right) as well as diagonally (up-left, up-right, down-left, down-right). Each movement incurs a small cost to induce an expectation of efficient action (by making longer action paths more costly), and reaching a food produces a reward.

The cost of action was sampled from an exponential distribution with *λ* = 100 (mean cost = 0.01) to ensure that results did not depend on any particular value of movement cost. This value determined the cost of moving in one of the four cardinal directions, with the cost of moving in any diagonal direction given by 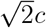 (by the Pythagorean theorem).

The reward was set to be large (*R* = 50) so that it would always outweigh the costs (i.e., to avoid situations where the food is not rewarding enough to justify the cost of getting it; as this has never been observed in the experimental paradigms under consideration). Note that exact reward values have important consequences in environments where different objects have different rewards, but this is not the case in our paradigms, where all food rewards were always identical.

MDPs further require that rewards are temporally discounted, such that an immediate reward is more valuable than a reward that can be obtained in the future. The temporal discount factor was set to slightly favor immediate rewards over future ones (*γ* = 0.95).

Equipped with this formulation, MDPs make it possible to calculate which actions an agent should take to obtain the highest possible reward (called a policy). Standard algorithms focus on obtaining the optimal solution, no matter the difficulty of the problem. In reality, as the value of going for two alternative food rewards become more similar, it should be harder to reliably identify which one is better (e.g., it is obvious that it’s better to get a reward that is one step away rather than an identical fifty steps away; but it would be harder differentiate if the rewards difference were, say, thirty-three and thirty-four steps away in different directions). To account for this, we made the MDP policy probabilistic, using the standard approach of softmaxing the value functions (Jara-Ettinger et al., 2020; Jern et al., 2017; Baker et al., 2017; Lucas et al., 2014) using a temperature parameter of *τ*_choice_ = 0.05. At this setting, in the Center-Wall paradigm (Fig. 2), the probability of the subject pursuing the center food (value 50, three steps away) vs. the food on their side of the barrier (value 50, four steps away) is about 0.51. This is in keeping with the spirit of the experiment, where small differences in physical distances are not considered main drivers of behavior (Hare et al., 2000); instead, behavior depends on the NHPs’ expectations of the relationship between seeing and knowing. To ensure our conclusions do not hinge on this particular value of the softmax parameter, we conducted robustness analyses. Increasing the value of the softmax parameter produced small quantitative changes in the exact model fits, but the key conclusions remained the same. Only extremely small values of the softmax parameter (which result in hypersensitivity to small differences in path lengths, describing an organism where even the smallest differences in cost produce massive differences in behavior) meaningfully affected the models’ performances. See SM Section 1.3 for details.

## Modeling competitor behavior

So far, our framework has specified (1) the general social behavior where the NHP avoids the reward they expect the dominant competitor will choose, (2) a tradeoff between egocentric and social behavior (governed by a reliance parameter), and (3) how egocentric behavior is determined. The last remaining component is how the NHP makes predictions about their competitor. We explored this through seven computational models, summarized in Table 2. While these models can be thought of as points in a continuum of representational complexity, they can be roughly organized into three broad categories of accounts: *Non-mentalistic, Semi-mentalistic*, and *Full Theory of Mind*. For simplicity, here we introduce the high-level intuition behind each model, and a formal presentation is available in SM Section 1.

**Table 2.** Summary of computational models. See SM Section 1 for technical presentation.

| Family | Model | Summary |
| --- | --- | --- |
| Non-mentalistic | Egocentric | NHP always acts egocentrically.<br>No social prediction mechanisms. |
|  | Food-directed | Competitor has an equal probability of going to any food in the room. |
|  | Location-directed | Competitor has an equal probability of going to any actual or possible food location |
| Semi-mentalistic | True belief | Assume shared knowledge. |
|  | Partial representation | Represent only what's in shared view. |
|  | Uncertain representation | Competitor knows about everything they see and there's 50/50 chance they know about any object they do not see. |
| Full ToM | Ignorance representation | Accurate representation of what the competitor knows and does not know. |

### Non-mentalistic models

The first family of accounts captures the idea that NHPs do not represent other minds. We started with the simplest account, the *egocentric* model, where NHPs entirely ignore the competitor. According to this model, NHPs simply pursue the food reward reachable by the shortest path (Fig. 1b), determined using the egocentric behavior model described in the general framework.

We next considered two accounts where NHPs predict behavior by relying on simple behavioral cues. These models are more sophisticated in that they require the subject to estimate how their competitor might behave. Thus, these models (and all models in more complex families) now also rely on MDPs to model how the competitor will move in space. While MDPs are often used to model first-person behavior, research has shown that they can also be used to model expectations about how other agents will act to obtain rewards, therefore serving as a framework for Theory of Mind (Baker et al., 2017; Jara-Ettinger et al., 2020; Baker et al., 2009). Critically, MDPs model behavior through an assumption that agents move efficiently toward their goals—an assumption that both human and non-human primates share (Gergely & Csibra, 2003; Rochat et al., 2008).

In the *Food-directed behavior* model, the subject expects the competitor to pursue one of the food rewards (possibly learned from experience seeing conspecifics go toward food) but lacks a mechanism for predicting which one, therefore placing an equal probability over each food reward (Fig. 1c), regardless of whether they are visible or not. In these paradigms, because there are always two food rewards, this model chooses each with 50% probability. This model can therefore be considered as a baseline representing chance performance.

The *Location-directed behavior* model extends behavioral expectations to include the expectation that competitors might also check locations that they cannot see (which could also be learned from experience observing conspecifics without necessarily representing epistemic states). This model therefore places equal probability over all food rewards and places where food could be located (even if none is there; Fig. 1d).

### Semi-mentalistic models

The *semi-mentalistic* family consists of three models that represent the competitor’s mind, but lack full human-like mental state representations. These models are loosely inspired by proposals that NHPs do represent other minds, but in a markedly limited way (e.g., Martin & Santos 2016).

The *True belief* model captures the idea that NHPs attribute their own knowledge to others and expect competitors to act based on this knowledge (Fig. 1e). Although the subject and competitor share the same representation of the world, they may still pursue different food rewards (e.g., each individual might go for their closest food reward, which could be different for the subject and the competitor).

In the *Partial representation* model, the NHP computes what parts of the environment are in the shared visual field and includes only those shared representations in their model of the conspecific’s mind. This is inspired by the idea that NHPs might be able to tag some of their own representations as shared with others, but cannot create representations for the conspecific that are different from their own (i.e., cannot represent the conspecific as not knowing something that the subject knows). This model therefore has no representation of the competitor’s awareness (or lack thereof) of objects or locations hidden from the competitor’s view (Fig. 1f). It also does not represent areas that the conspecific might be able to see, but the subject cannot. That is, any objects outside of the shared visual field are simply not represented. This model therefore creates a simplified representation of the environment that is limited to what’s in the shared visual fields, and then predicts how the competitor would act in that shared visual space.

In the *Uncertain representation* model, the NHP assumes the competitor is aware of any object in their visual field, but is uncertain whether the competitor is also aware of objects outside their field of view. This model differs from the *Partial representation* model in that the subject considers the possibility that the conspecific might know about the hidden reward (but is unsure whether they know about it or not). This model therefore builds predictions by integrating the two epistemic hypotheses about the competitor (knows or does not know about the hidden object, using a uniform prior; Fig. 1g) for every possible location that is not in their visual field.

## Full Theory of Mind model

Finally, the *Full ToM* family consists of theories of human-like Theory of Mind. Although there are a variety of different proposed theories of what full human-like ToM consists of (e.g., Hutto 2012; Gordon 1992; Jara-Ettinger 2019), these theories only make different predictions in complex cases that go beyond those captured in the paradigms considered here. So for this family, we include only an *Ignorance representation* model, where NHPs have an accurate representation of their competitor’s visual perspective, representing both their ignorance of hidden objects and their knowledge of visible objects (Fig. 1h).

## Results

Our analysis consisted of three stages. First, we looked at the qualitative performance of our models before integrating reliance. We next examined the quantitative performance to compare predicted to observed effect sizes. Finally, we added the reliance parameter and fit it to empirical data to reveal the relation between posited representations and inferred reliance.

### Qualitative model performance

To understand each model’s behavior, we began by evaluating its capacity to replicate the qualitative pattern of successes and failures documented in the NHP literature, assuming full reliance (i.e., that primates always use their expectations about the competitor to guide their behavior; *r* = 1). This allows us to understand each model’s core predictions and see which accounts can capture the broad qualitative patterns of NHP behavior. For this analysis, each experiment was coded as showing a directional effect or not based on results reported in the literature (i.e., whether the results were significantly above chance or not). This yields directional effects for the *Center-Wall, Open-Hidden*, and *Transparent-Hidden* paradigms and chance on the *Hidden-Hidden* and the *Open-Transparent* paradigms (Figure 2).

To compare model predictions to these binary outcomes, we need to convert each model’s continuous prediction of effect size into a binary decision. Simply checking whether the model’s prediction differs from chance (i.e., 0.5) would misclassify small effects as directional—effects that would never produce a significant result in NHP studies given typical sample sizes. To avoid this issue, we reviewed the sample sizes used in the NHP studies and estimated the smallest effect size that would produce a significant result in those experiments. We then used this threshold, 0.6, to classify model predictions (see SM Section 4.2 for details).

Fig. 3 shows each model’s ability to replicate the qualitative NHP results for each paradigm. This revealed that the non-mentalistic models were outright unable to capture even the qualitative pattern of results, failing to make a directional prediction on the three paradigms on which NHPs succeed (Center-Wall, Open-Hidden, and Transparent-Hidden Routes). Only the full model (*Ignorance rep.*) and two of the three semi-mentalistic models (*Uncertainty rep.* and *Partial rep*, but not *True Belief*) replicated the qualitative experimental record.

**Figure 3.**
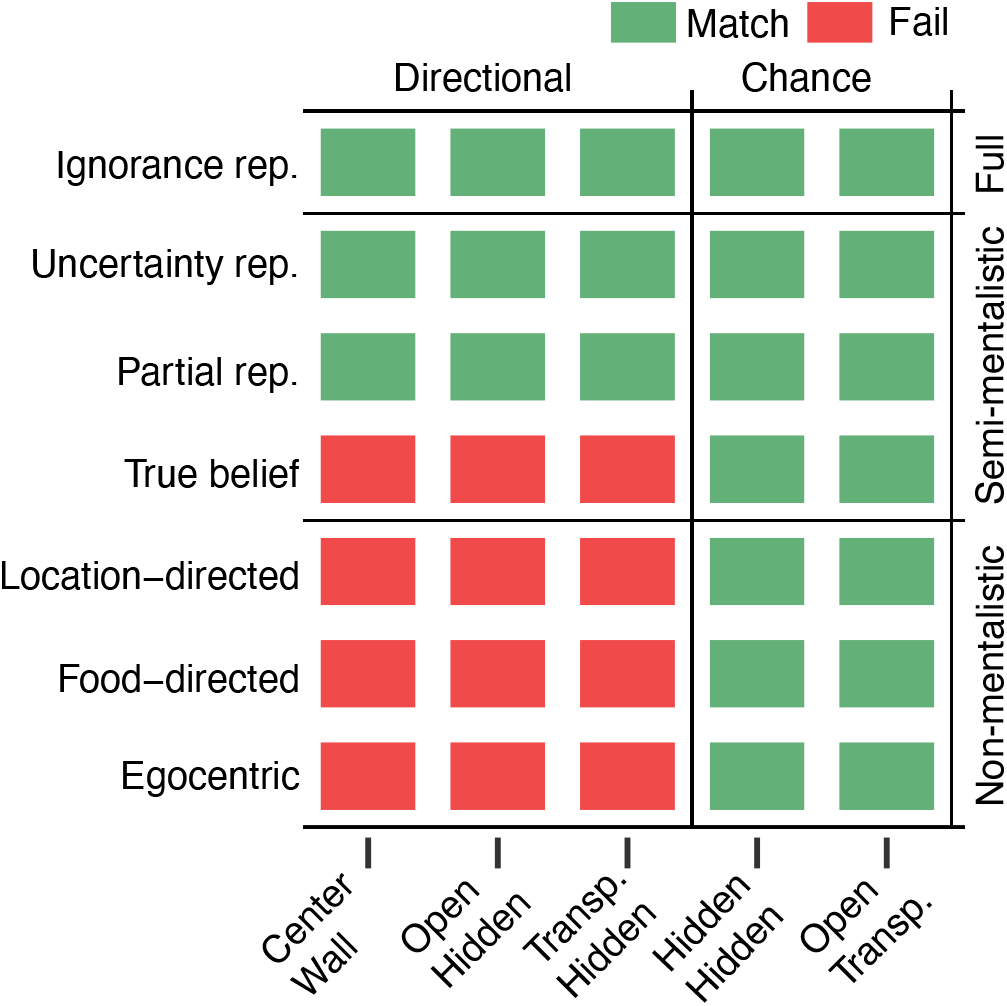
Results from qualitative model evaluation assuming full reliance. Each row represents a computational model, and each column represents one of the five paradigms. NHPs show preferences for one food location over the other on the first three directional paradigms and equal preference on the last two control paradigms. Color indicates whether the model’s prediction is consistent with NHP behavior or not, using a threshold of t = 0.6 to differentiate predictions of a directional vs. chance effect.

These qualitative results align with current debates surrounding NHP perspective-taking and Theory of Mind, lending validity to our core computational framework. Theories where NHPs predict behavior by relying on learned associations and past experience (similar to those in the *non-mentalistic* family of models) have largely fallen out of favor in these broader debates, while current debates focus on whether NHPs have a human-like ToM or a simpler mentalistic representation (similar to those in the *semi-mentalistic* and *Full ToM* families; e.g., Martin & Santos, 2016; Krupenye & Call, 2019).

### Quantitative model performance

Having found that several models could replicate the qualitative pattern of results, we next analyzed the exact effect sizes that each model predicts. This allows us to reveal whether a model, even when it makes the correct directional prediction, may lack explanatory power by either over- or under-predicting an effect size relative to observed NHP behavior. Therefore, we next directly compared the predicted and empirical effect sizes for each paradigm. To make these effects easier to interpret, we also replicated these paradigms with human participants in a simple online game, obtaining adult human effect sizes as a reference point. The full experimental procedure for the human experiment is available in SM Section 2.

Fig. 4 shows each model’s performance across the five paradigms, overlaid with human and NHP results^1^. A detailed walk-through of each model’s predictions is available in SM Section 3. In each plot, black lines represent model predictions and shaded regions represent empirical data, such that a closer fit between the black line and the shaded region indicates better model-data alignment.

**Figure 4.**
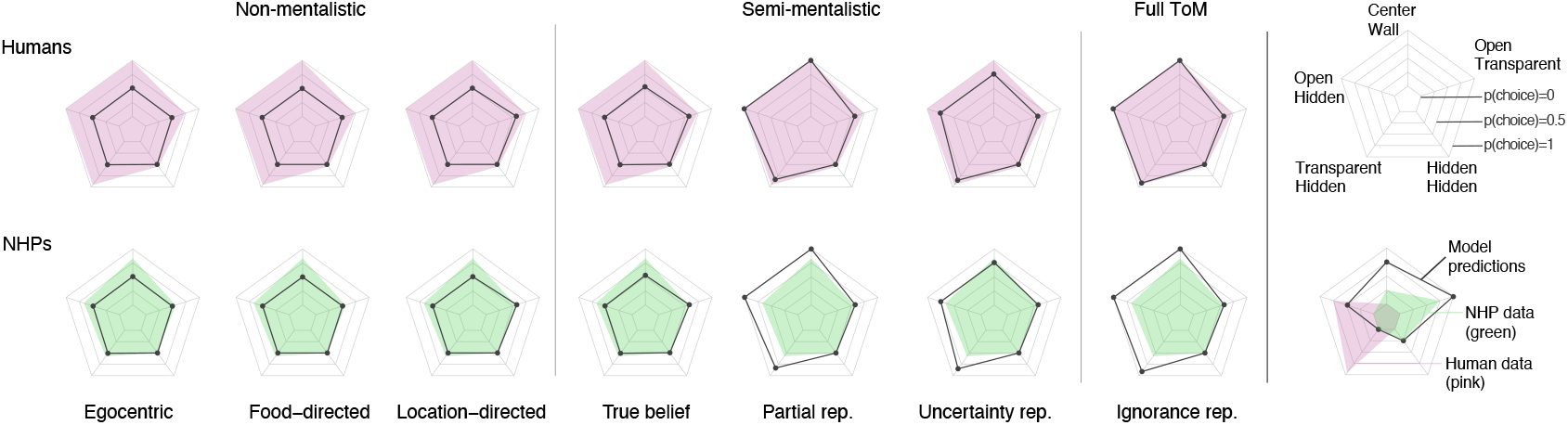
Quantitative model predictions assuming full reliance. Each radar plot shows model-predicted effect sizes (black lines) compared to empirical data (human in pink, top row; NHP in green, bottom row), for each of the five paradigms (one per corner). Values represent the probability of choosing hidden food for the directional paradigms and the probability of choosing right-most food in control paradigms. A tighter fit of the black line to the colored region indicates better model-data alignment.

For human data (top row, pink), all non-mentalistic models under-predict effect sizes (i.e., the pink region exceeds the black contour), and so did the *true belief* model in the semi-mentalistic family. In contrast, the other two semi-mentalistic models and the full ToM model all showed a closer alignment with human behavior.

For NHP data (bottom row, green) a different pattern emerged. Here, the four simplest models (the non-mentalistic models and the *true belief* model) under-predicted the observed effects, while the three most complex models over-predicted the effects in some paradigms.

To confirm this visual assessment quantitatively, we would ideally calculate each model’s explanatory power over the empirical data, *P* (*D|M*). However, the trial-level performance data needed for this analysis is not available for every paradigm (see SM Section 4.1 for details), so we begin by computing the root mean squared error (RMSE) between model predictions and empirical data, allowing us to include all paradigms. This analysis mimics the visual assessment by weighting each paradigm equally, regardless of how much data is available per paradigm. Additional information and full results are available in SM Section 3.2.1.

The RMSE analysis confirmed the visual inspection of results. Compared to humans, the non-mentalistic models produced the largest errors (RMSE*>* 0.376 for all cases). Among the semi-mentalistic models, the *True belief* model provided an error comparable to the non-mentalistic models (RMSE= 0.367), the *Uncertain rep.* model’s error was about half that of the *True belief* model (RMSE= 0.174), and the partial representation had again about half the error of the *Uncertain rep.* (RMSE= 0.092) model. Finally, the human ToM model (*Ignorance rep.*) showed the lowest error (RMSE= 0.077).

Comparing the models to NHP data showed a different pattern of results. The largest discrepancy was against the human-like ToM model (*Ignorance representation*; RMSE= 0.225), followed by the *Partial rep.* model (RMSE= 0.203). The next three worst fits were the models in the non-mentalistic families (all RMSE*>* 0.167). Finally, the best two models were in the semi-mentalistic family: *True belief* (RMSE= 0.156) and *Uncertain rep.* (RMSE= 0.134).

Together, these results suggest that the *Ignorance representation* model best fits human behavior, whereas the semi-mentalistic *True belief* and *Uncertain rep.* models best fit NHP behavior. This is because the non-mentalistic models do not capture NHP behavior, and the human-like full ToM model predicts effect sizes that are much stronger than those documented.

### According to each model, how often do NHPs rely on their social representations?

Our analyses so far have focused on understanding each model’s core performance, which clarifies how we would expect NHPs (or humans) to behave if every individual always used these representations in every experiment. Having characterized how these models perform, we are ready for our key analysis of interest: formally estimating how much NHPs *rely* on their social representations, according to different theories. Our goal is to reveal the implicit assumptions that each account carries about how frequently its representations are used for the account to explain the data. This assumption is usually left unstated, but here, our computational framework allows us to extract it. Specifically, we accomplish this by searching over the space of possible reliance values and finding the one that maximizes each model’s potential to explain NHP data. This approach uncovers each theory’s implicit commitment to how frequently NHPs rely on their social representations to guide their behavior.

To find the value of reliance that best explains behavior, it is necessary to look beyond point estimates of the effect sizes and instead consider the full distribution of trial-level successes and failures. We approached this by finding the reliance parameter that maximizes each model’s probability of generating the data, *P* (*D|M*). Unfortunately, choice data were not always recoverable in the eleven NHP experiments we considered (Table 1), forcing us to limit our analysis to seven experiments and to drop the hidden-hidden paradigm. See SM Section 4.1 for details. We then calculated the probability of each model generating the behavioral choice data (i.e., the model’s explanatory power), testing reliance values between 0 and 1 by intervals of 0.01 (see SM Figure S2). We conducted this analysis twice, once fitting reliance to explain human data, and another fitting reliance to explain NHP data.

We begin by evaluating the model fits against human judgments to confirm the validity of our analysis. Here, given the task’s simplicity, we should expect that the best fit model is the human-like ToM one with a high reliance parameter. This was confirmed by our analysis (Fig. 5A). The best model for explaining human data was the *Ignorance rep.* model, with a posited *r* = 0.99 reliance.

**Figure 5.**
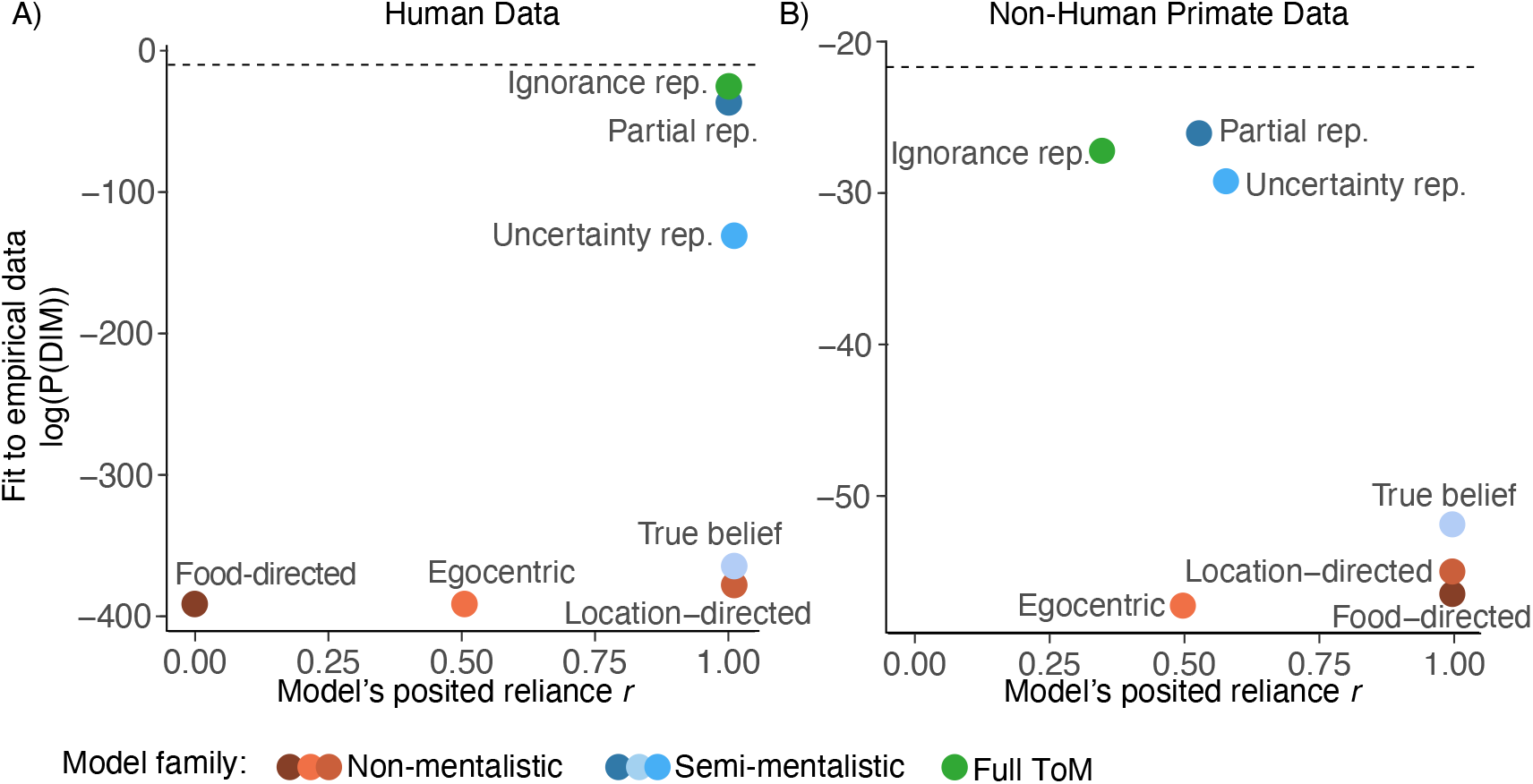
Fitting each model’s reliance to explain the empirical data for humans (panel A) and non-human primates (panel B). Each plot shows the reliance parameter (r; x-axis) that maximizes explanatory power (y-axis, in log-space). Thus, models with higher values on the y-axis indicate better explanatory power, whereas models with higher values on the x-axis indicate that this explanatory power is accomplished by assuming most subjects are using the posited representations. The absolute values on the y-axes differ because the two datasets have different amounts of data. For each dataset, the maximum possible explanatory power a model could achieve (a ‘ceiling’) is represented by a dashed horizontal line.

Among the semi-mentalistic family, all three models maximize their fit to human data by positing high degrees of reliance (*r* ≥ 0.99 in all cases), but they do not capture human data as well as the *Ignorance rep.* model. The *Partial rep.* was the closest (Partial: *LL* = −36.3, Ignorance: *LL* = −25.2). By contrast, the *Uncertainty rep.* and *True belief* models provided substantially lower fits (Uncertainty: *LL* = −130.9, True belief: *LL* = −364.5), showing that it is, in principle, possible for these paradigms to distinguish between subtly different accounts. Finally, the models in the non-mentalistic family all showed the lowest explanatory power. Indeed, the *Food-directed* model was such a poor fit to human judgments that its best explanatory power was reached under reliance *r* = 0, revealing that even the smallest use of this model negatively affected model fit.

To better interpret how well the models explain the data, we also calculated the explanatory power of the best possible model (a theoretical ‘ceiling’ model that perfectly predicts the exact observed choice probabilities for each paradigm). This model reached *LL* = −10.0, providing a ceiling on explanatory power (depicted in Fig. 5A by the horizontal dotted line), or a measurement of how difficult this dataset is to explain. Comparing the explanatory power of the *Ignorance rep.* model with this ceiling (Δ*LL* = −15.2) reveals room for improvement (discussed in SM Section 2.5).

Having validated our approach against human data, we next applied the same approach to reveal the implicit reliance each theory must posit to explain NHP behavior. The results are shown in Fig. 5B. Consistent with the qualitative analyses, models in the non-mentalistic family showed the worst fit to NHP data. This suggests that these models are outright unable to explain NHP behavior, even when they are allowed to posit different degrees of reliance.

The models in the semi-mentalistic family showed variable performance. The *True belief* model had explanatory power nearly as low as the non-mentalistic models. By contrast, the *Partial rep.* and *Uncertainty rep.* were both able to achieve among the highest explanatory power of the exact effect sizes (*LL* = −26.1 and −29.2, respectively). To do so, however, required positing different degrees of reliance (*r* = 0.53 and *r* = 0.58, respectively).

The human-like *Ignorance rep.* model also reached among the highest explanatory power (*LL* = −27.2), but to do so, it required positing a surprisingly low reliance: *r* = 0.35. This is because this model predicts effect sizes that are much stronger than those found in NHP behavior.

This analysis reveals a previously hidden assumption in the debate over whether NHPs have a human-like ToM or not: if NHPs are using human-like representations in these tasks, then they must only use them about 35% of the time. Alternatively, if NHPs are using a simpler type of ToM, then they must only use it about ≈ 55% of the time.

Finally, comparing the explanatory power of these three best models to the ceiling of how well this dataset could, in theory, be explained (*LL* = −21.7) shows that these models still leave some room for improvement (*Partial rep.*: Δ*LL* = −4.4; *Uncertainty rep.*: Δ*LL* = −7.5; and *Ignorance rep.*: Δ*LL* = −5.5). This leaves open the possibility that other mental representations may better explain NHP behavior on visual perspective-taking tasks.

### General Discussion

Here, we introduced a computational framework for comparative social cognition and applied it to visual perspective-taking as a case study. First, we implemented possible theories of non-human primate (NHP) social representations as computational models and tested them on five paradigms from the perspective-taking literature. Second, we formalized the notion of *reliance* to explicitly model the frequency with which NHPs would have to use their posited representations in order for each account to explain the effect sizes observed in the NHP literature. Using these two key components, our framework allowed us to find the combinations of representations and reliance that best explain NHP behavior on classical visual perspective-taking tasks.

Our paper produced three broad sets of results. First, we found that a variety of mentalistic theories are able to explain the qualitative pattern of NHP results equally well (Fig. 3). This finding provides a computational replication of the sociological state of the field, where there is debate about whether NHPs have a human-like ToM or not, but broader agreement that they have some form of mentalistic representation (e.g., Martin & Santos, 2016; Krupenye & Call, 2019).

Second, we used our computational models to derive the effect sizes that each theory predicts. This revealed that the models with the most complex representations best fit human behavior, while models with simpler representations of other minds best fit NHP behavior.

Finally, by considering the possibility that NHPs (and humans) don’t always rely on their social representations, we fit the degree of reliance for each model. This reliance parameter allowed us to estimate how frequently subjects would need to use the posited representation in order for a given account to explain the data. For example, the account that best fit human perspective-taking data is one where people use a complete Theory of Mind almost all the time. In contrast, capturing NHP behavior required reducing reliance in all models. Specifically, the most human-like model of Theory of Mind could only explain primate behavior by assuming that NHPs use these representations about one-third of the time. Therefore, positing that NHPs have human-like Theory of Mind representations comes with the implication that NHPs only rarely use those representations. We also found that two other mentalistic accounts, which posit that NHPs use simplified representations of other minds (*Partial representation* and *Uncertain representation*; Table 2), can capture the data, but only if their reliance is just over 50%.

Taken together, this approach converges with past investigations to validate the mentalistic nature of NHP visual perspective-taking, while also revealing a key difference between human and non-human primate social representations: the degree to which they are relied upon to guide behavior. This trade-off between representational complexity and reliance is especially pertinent to questions surrounding whether NHPs have a human-like Theory of Mind. Our work suggests that, if NHPs have a Theory of Mind of similar representational complexity to humans, then its usage is surprisingly limited compared to how readily human adults use their Theory of Mind, at least in visual perspective-taking tasks. This is particularly interesting given that the experimental paradigms considered in this paper were designed for ecological validity, using competitive situations with known conspecifics (Hare & Tomasello, 2004). By contrast, human Theory of Mind is characterized not only by its cognitive richness but also by how often we use it in a wide range of contexts, even automatically in some cases (Kampis & Southgate, 2020). For instance, people will sometimes automatically track each other’s knowledge from their visual perspective (Samson et al., 2010) and make automatic common ground inferences (Rubio-Fernández et al., 2019). One further question, however, is whether human reliance on Theory of Mind would decrease if participants were tested under stressful, time-constrained conditions that more closely approximate the competition paradigms used with NHPs. In such cases, humans may switch to non-mentalistic strategies to predict the behavior of other agents (Andrews et al., 2021). We hope that our work enables future discussions of comparative Theory of Mind to achieve more nuance by incorporating claims related to both representations and reliance.

Our work has three primary limitations. The first is that we only focused on seven computational models of visual perspective-taking. While these models varied substantially in their degree of cognitive complexity, they do not cover the full space of conceptual theories. We hope that in future work, researchers developing theories of NHP social behavior can work with modelers to implement their theories. Our framework then provides a standardized way to estimate how well a theory captures NHP data and how often NHPs rely on the representation in order for the theory to hold. From this standpoint, our computational models can serve as a set of benchmarks that help quickly reveal whether new theories have strong explanatory power.

The second limitation of our work is that formalizing theories into computational models required us to set multiple parameters: the cost of movement, the magnitude of the food reward, the future discount, and the softmax parameter. Here we took a conservative approach: minimizing their influence when possible (as with future discount and food reward), integrating over plausible values when unsure (as with costs), and conducting robustness analyses when their influence was likely to matter (as with the softmax parameter). While we did not find that parameter choices produce overwhelming differences, they inevitably affect the exact predicted effect sizes and the estimated reliance parameters. This is not a flaw. It instead reveals that factors often overlooked in theories of NHP social reasoning might be critical for theory evaluation. When parameters in a model affect predictions, this is not a weakness of a computational model, but rather evidence that empirical work should attend to and study these components. We hope future research can estimate baseline energy expenditure, food motivation, and future discount parameters in tasks so as to guide parameter settings and even account for individual variation.

One final limitation of our work is that we exclusively focused on studies conducted with catarrhine primates: chimpanzees and Tonkean macaques. We chose this level/unit of analysis for two reasons: first the catarrhine group includes humans, thus making other catarrhine species most relevant human-nonhuman primate comparisons and, second, catarrhines typically perform well on visual perspective-taking tasks and are often considered to not vary in this capacity, making them easier to model/evaluate as a group (Lewis & Krupenye, 2022). Our modeling approach can easily be applied to a wider range of species in the future as more data becomes available since perspective-taking capacities could vary dramatically across the primate taxon. Indeed, initial evidence suggests that capuchin monkeys (a species of platyrrhine, more distantly related to humans relative to catarrhines) and ring-tailed lemurs (a species of strepsirrhine, even more distantly related to humans relative to catarrhines and platyrrhines) are more limited in their understanding of the relationship between perception and knowledge relative to chimpanzees and macaques (Bray et al., 2014; Hare et al., 2003). As such, the computational models that were discarded here for their poor qualitative fit to the catarrhine data (e.g., *Location-directed* and *True Belief*) might prove more useful for characterizing the social cognitive representations of platyrrhines and strepsirrhines. Similarly, as more data become available for additional catarrhine species, our approach could be used to produce an internal species-level analysis detect nuanced internal distinctions within catarrhines, providing further insight into the evolutionary origins of primate Theory of Mind.

While our focus was on visual perspective-taking tasks, our general framework can be adapted to Theory of Mind’s many other components, including gaze-following (Tomasello et al., 1998; MacLean & Hare, 2012; Rosati et al., 2016; Drayton & Santos, 2017; Bettle & Rosati, 2019), awareness representations based on an agent’s past perceptions (Hare, 2001; Santos et al., 2006; Drayton & Santos, 2018; Horschler et al., 2019, 2021), and belief representations (Kaminski et al., 2008; Krupenye et al., 2016; Martin & Santos, 2016; Kano et al., 2020; Horschler et al., 2020a,b). Our work offers a general methodological approach that shows how experimental paradigms can be standardized and placed in a common representational framework that allows for the design and evaluation of computational models.

Extending our framework to other Theory of Mind abilities, however, will require additional attention to the construct of reliance. In our current framework, reliance is a broad construct that captures many forces that may shape performance, such as motivation, the “effort” required to deploy representations, and cognitive load (e.g., time pressure, uncertainty, etc). While our current framework subsumes all these factors under the construct of reliance, we hope that future work can use this construct to test different causes behind degrees of reliance. For example, future versions could extend this approach to capture paradigm-specific reliance (i.e., some experimental paradigms might increase NHPs’ reliance on their available social representations; e.g., Krupenye et al. (2016)) and possibly even model switching (i.e., NHPs might use different representations in different tasks). Even within visual perspective-taking tasks, some variations in the paradigms appear to reduce visual perspective-taking success (Povinelli et al., 1996). These failures have been interpreted as revealing that NHP social cognition is most visible in ecologically-relevant competitive contexts (Hare, 2001) and where food rewards are far enough apart that they enhance attention to physical costs. These hypotheses about the basis of paradigm-specific differences in NHP behavior can be quantitatively tested using our reliance construct. Thus, our theory-testing framework can help future researchers test not only the representational underpinnings of a given capacity but also the factors that impact specific task performance.

## Conclusion

Understanding what non-human primates (NHPs) know about each other’s minds has been a central goal in comparative cognitive science for over forty years. Progress has been challenging, however, because NHPs can fail to reveal their capacities in some tasks or succeed through simpler strategies in others.

Our work advances this debate by computationally formalizing the predictions of different theories and estimating how each theory must posit different degrees of reliance to account for behavior. Our results show that many simple, non-mentalistic theories of NHP social behavior outright fail. Yet, both human-like and non-human-like representations can explain NHP behaviors, but each assumes different degrees of reliance on these representations. This means that, regardless of which theory one favors, NHPs do not resemble humans in how frequently and pervasively we use our Theory of Mind. Perhaps NHPs exist in social environments where applying Theory of Mind is not often necessary, possibly because they have alternative role-based systems for understanding behavior (Jara-Ettinger & Dunham, 2024). But this does not mean they lack mentalistic representations; on the contrary, our framework shows strong support for them. Thus, rather than being deflationary, our findings invite researchers to explore when NHPs use their Theory of Mind and when they rely on alternative non-mentalistic computations to solve complex social problems.

## Acknowledgments

We thank Mel Andrews and Max Siegel for helpful comments.

## Footnotes

1 For the Hidden-Hidden paradigm, the NHP study reported a null result, but did not report the exact effect size (Hare et al., 2000), so we set it to 50%.

