## Supplemental Materials for "What Primates Know About Other Minds and When They Use It: A Computational Approach to Comparative Theory of Mind"

### Supplemental Material

---

---

#### 1. Implementation of computational models

The main paper details the high-level computational structure of the model and each of the seven computational models for predicting conspecific behavior. This section provides additional information as to how each model was implemented.

Each of the five paradigms (Fig. 2 in Main) are represented as a simple gridworld. In all simulations, the competitor always started from the same initial location in the gridworld, and the subject always started from the same initial location opposite the competitor. Food rewards and barriers were set to match the key variables of the experimental paradigm (e.g., equidistant from subject, etc). All five map descriptions are available in our OSF repository: https://osf.io/qjn9m/?view_only=a8ede1cffb684452af9b2c54cb5ac15c.

In all cases where we use MDPs, behavior was estimated via Monte Carlo sampling, using 5000 samples per MDP, such that an MDP's probability of choosing a food item equals the proportion of times that this reward was chosen in the simulations. Final model predictions were then obtained by combining the probabilities generated from different MDPs as specified by the model description (e.g., averaging over the MDPs in cases where the subject holds multiple hypotheses).

##### 1.1. Implementation details specific to the Transparent-Hidden Routes paradigm

In most paradigms, we used MDPs to model how the subject represents the competitor's movements toward the food rewards. The one exception was the Transparent-Hidden routes paradigm. In this paradigm, the subject considers how likely the competitor is to detect them along each of two routes to the foods. Therefore, the MDP represents how the competitor reasons about the subject's movements (rather than how the competitor moves towards food rewards). For each action plan (i.e., route the subject could take), the model then integrates a probability that the competitor would detect the subject, and the probability of obtaining a reward is the probability of completing the route undetected. For most accounts, the probability of detection along the transparent route was sampled from a distribution to account for the possibility that the competitor might not always notice the subject's movement or react on time, while also ensuring that model predictions do not depend on any particular value of the detection probability. Detection probabilities for each simulation were sampled from a Beta distribution with $\alpha = 9$ and $\beta = 1$ (giving a mean of $\mu = 0.9$).

##### 1.2. Details of each model implementation

###### 1.2.1. Physical planner

The physical planner simply executed the egocentric behavior MDP detailed in the main paper. This is equivalent to setting the reliance parameter $r = 0$, such that any predictions of the competitor are irrelevant (and hence not even specified).

###### 1.2.2. Food-directed behavior

To implement this model, each paradigm map was initialized as two MDPs, each with the agent in the competitor's starting position. Only one of the food rewards was present in each MDP, but all barriers were present in both MDPs. Together, these MDPs therefore generated the expected behavior of the competitor if it were going to each food reward, and predictions were subsequently averaged across MDPs (with equal weighting). This approach modeled an expectation that the competitor would choose one of the food rewards but lacked a mechanism for predicting which option the competitor may favor, therefore placing an equal probability on each location or route. Because this model specifies a behavioral expectation that competitors pursue food rewards (but lacks a mechanism for reasoning about how they detect others), the probability of detection in the Transparent-Hidden paradigm was set to $p = 0$.

###### 1.2.3. Location-directed behavior

To implement this model, each paradigm map was initialized similarly as the *Food-directed behavior* model, but also included MDPs with a hypothetical food reward in each location hidden from the competitor (e.g., behind the wall that never actually hid a food reward from the competitor in the Center-Wall paradigm). This approach therefore modeled an expectation that the competitor may search by considering all possible combinations of the locations where the competitor might see or look for food (including areas behind barriers where no food is currently present). Because this model expresses the idea of learned behavioral patterns, in the Transparent-Hidden Routes we apply this principle to detections, such that the subject has general expectations that their behavior is sometimes detected and sometimes not. Therefore, this expectation applied to the detection probabilities assigned to each route rather than to the food reward locations (i.e., considering all possible combinations of detection probabilities $p \sim Beta(\alpha = 9, \beta = 1)$ and $p = 0$ for transparent and opaque routes).

###### 1.2.4. True belief

To implement this model, each paradigm map was initialized as a single MDP with the agent in the competitor's starting position. All food rewards and barriers were present on the map. This approach therefore modeled a true belief default, where the subject attributes their own knowledge to competitors, and uses this representation to predict their behavior (note that this representation does not imply that both agents will pursue the same reward; e.g., each agent might go for the food reward closest to them). In the Transparent-Hidden Routes paradigm, detection probabilities for both routes were set to $p = 1$, following the idea that the subject always represented that the competitor shared the subject's knowledge (including their location).

###### 1.2.5. Partial representation

To implement this model, each paradigm map was initialized as a single MDP with the agent in the competitor's starting position. However, this MDP excluded areas of the map, barriers, and food rewards from the parts of the paradigm map that were occluded from the competitor's view (essentially cutting out pieces of the map; map files available in the OSF repo). This approach therefore modeled the possibility that the subject was entirely unable to represent the competitor's awareness of objects or locations hidden from their view, and instead used a limited representation that only considered regions that were visible to both agents. In the Transparent-Hidden Routes paradigm, the transparent route had a detection probability $p \sim Beta(\alpha = 9, \beta = 1)$, since the subject expects the competitor to share information in common ground. The hidden route was not represented in the map and therefore had no associated detection probability.

##### *1.2.6. Uncertainty representation*

To implement this model, each paradigm map was initialized as MDP(s) with the agent in the competitor's starting position. In each MDP, all food rewards that were visible to both the competitor and subject were included on the map, as were all barriers. In paradigms that included food rewards visible to the subject but not the competitor, we included one MDP with these food rewards present on the map and another MDP with them absent from the map, and predictions were subsequently averaged across MDPs. In the Transparent-Hidden Routes paradigm, this approach applied to the detection probabilities assigned to each route, such that the detection probability applied to the hidden route integrated over the two hypotheses about detection rate ($p = 0$ and $p \sim Beta(\alpha = 9, \beta = 1)$) using a uniform prior, resulting in detection probabilities sampled from $p \sim \frac{Beta(\alpha=9,\beta=1)}{2}$. The detection rate for the transparent route was sampled as $p \sim Beta(\alpha = 9, \beta = 1)$. This approach therefore modeled the possibility that the subject assumed common knowledge of objects and routes that were in plain sight for the subject and competitor, but had uncertainty about whether the competitor could see movement along the hidden route.

##### *1.2.7. Ignorance representations*

To implement this model, each paradigm map was initialized as a single MDP with the agent in the competitor's starting position. All barriers were present on the map, but only food rewards visible to the competitor were present. In the Transparent-Hidden Routes paradigm, the detection probability applied to hidden routes was $p = 0$, while the detection probability applied to transparent routes was $p \sim Beta(\alpha = 9, \beta = 1)$. This approach therefore modeled human-like ToM in these tasks, such that the subject has a complete representation of the competitor's knowledge and ignorance based on the competitor's visual perspective and uses it to predict their actions accordingly.

#### *1.3. Additional model parameters and sensitivity analysis*

The MDPs contain additional parameters. The movement cost together with the temperature parameter of the softmax, $\tau_{\text{choice}}$, determines the models' sensitivity to any difference in path length to one food or another. This is best shown on the Center-Wall paradigm, where the food in the center was marginally closer to the subject than the food behind the wall. The *Physical planner* prefers the closer food, and the strength of that preference is determined by the combination of the movement cost and temperature parameter. Costs were sampled from an exponential distribution with mean $\lambda = 0.01$, ensuring that the results do not depend on any single value for movement cost. The results presented in main use the softmax parameter $\tau_{\text{choice}} = 0.05$. Under this setting, the distances have minimal impact on the models' behavior, so that the models behave according to their posited relationship between seeing and knowing. At $\tau_{\text{choice}} = 0.05$, the *Physical planner* model displays minimal preference for the closer food (P(center)= 0.51). Since the strength of this preference is sensitive to $\tau_{\text{choice}}$, we conducted a sensitivity analysis to ensure that our overall results to not hinge upon the particular value of $\tau_{\text{choice}}$ used.

Figure S1 shows the results of a sensitivity analysis varying $\tau_{\text{choice}}$ from 0.01 to 0.25. For each value of $\tau_{\text{choice}}$, we found each model's posited level of reliance that maximized its ability to explain the empirical NHP behavioral data. Figure S1A shows how each model's ability to explain the empirical NHP behavioral data varies as a function of $\tau_{\text{choice}}$.

When $\tau_{\text{choice}}$ is small, the models are highly sensitive to movement costs and act more efficiently to minimize them. At extremely small values of $\tau$, like $\tau = 0.01$, all models perform poorly. The *True Belief* model, incidentally, benefits from small values of $\tau$ in the range [0.01-0.03], achieving maximum explanatory power $LL = -25.6$, because these values allow it to better capture NHP directional behavior on the Center-Wall paradigm by assuming the competitor pursues the food in the open because it is closer. The same reasoning (about the physical cost of navigating around a barrier) also allows the *True Belief* model to capture the Open-Hidden paradigm. However, *True Belief* cannot, with any value of $\tau$, capture behavior on the Transparent-Hidden Routes paradigm because both paths are equally costly.

As $\tau_{\text{choice}}$ increases to 0.05, the *Ignorance rep.*, *Partial rep.*, and *Uncertainty rep.* models perform better. As the models become less sensitive to small differences in movement cost, their behavior is driven more by what the competitor can and cannot see, and these models better match NHP behavior. The *Partial rep.* achieves the best fit to NHP behavior at 0.05, with $LL = -26.1$. *Ignorance rep.* explains the data slightly less well ($LL = -27.2$) and

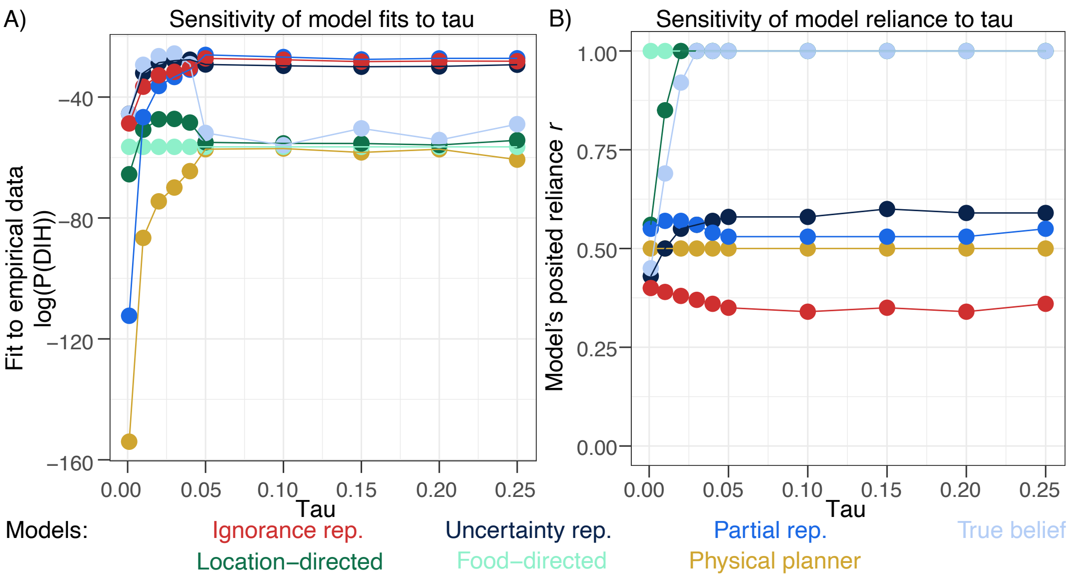

Figure S1: The results of a sensitivity analysis varying $\tau_{\text{choice}}$ (x axis). A) Each model's explanatory power (y axis) as a function of $\tau_{\text{choice}}$. B) The level of reliance (y axis) that each model posits to best explain NHP data as a function of $\tau_{\text{choice}}$. For $\tau_{\text{choice}} \geq 0.05$, results are qualitatively consistent. Values of $\tau_{\text{choice}} \leq 0.04$ create higher sensitivity to small differences in path lengths, enabling the *True Belief* model to perform best.

*Uncertainty rep.* does best at $\tau_{\text{choice}} = 0.04$, where it explains the data with ($LL = -27.6$). These three models' fits to NHP data remains qualitatively consistent for $\tau_{\text{choice}} = [0.05 - 0.25]$. For values of $\tau_{\text{choice}}$ in this range (Fig. S1B), each model's posited reliance also remains consistent: the Ignorance Rep model has to posit a small degree of reliance ($r \approx 0.35$) to best capture NHP behavior, while the two leading mentalistic models, *Partial Rep* and *Uncertainty Rep*, posit $r \approx 0.58$ and $r \approx 0.53$, respectively. Because the values of $\tau_{\text{choice}} \geq 0.05$ produce model behavior that reflects expectations of what the competitor can and cannot see rather than small differences in path length, and because these results are stable, the main text presents the results using $\tau_{\text{choice}} = 0.05$.

Because $\tau_{choice}$ modulates the MDPs' sensitivity to small differences in the expected reward for pursuing one goal versus another, the varying $\tau_{choice}$ produces similar effects as would varying expected rewards. Assuming that the rewards are the same for both foods, and that the cost of movement never outweighs the reward of obtaining the food (in which case the agent would not approach either reward), varying $\tau_{choice}$ produces the same effects as varying the rewards or costs.

Finally, the MDPs also considered the possibility that an agent might not always select the best action (step in the right direction). This was captured by another softmax with temperature parameter $\tau_{action} = 0.01$. This affected the exact paths that the agent took to a particular food, but not the choice of which food to pursue. Since the model predictions are always used in terms of the choice of food that the agent pursued (rather than the exact shape of the path), this parameter did not influence the model predictions relevant to any qualitative comparisons or analyses.

#### 2. Human experiment

To validate our approach, we ran a pre-registered experiment (see OSF repo) that tested adult humans on tasks conceptually equivalent to the classical NHP perspective-taking paradigms. This experiment was subject to an IRB exemption.

While we know that humans can succeed on some perspective-taking tasks from infancy (e.g., Luo & Johnson 2009), running conceptually equivalent tasks with adults online allowed us to both obtain performance that was more directly comparable to NHPs' and also allowed us to estimate human representational reliance. In NHP version of these tasks, the competitor is a dominant conspecific, which will behave aggressively towards the subordinate subjects, particularly in competitive food contexts. Our human task therefore framed the competitor as a 'bully' to maintain the sense of a dominance hierarchy and potential risk of retribution.

##### 2.1. Participants

200 U.S. participants (Age: mean=37.3 years, range=19-77 years) completed the task on Prolific. There were no exclusions.

##### 2.2. Stimuli

The stimuli consisted of maps of five grid worlds representing the five perspective-taking paradigms. The maps showed transparent and opaque barriers and markers indicating the participant's and the competitor's locations.

##### 2.3. Procedure

Human participants were introduced to a candy-gathering task in which they competed against a bully. They were told that the bully would always go for one of two identical candies. If the bully and the participant pursued the same candy, then the bully would steal it. But if the bully and the participant chose different candies, they would both get to eat their candy in peace. Participants viewed the five grid world maps in randomized order and clicked on the candy they would choose to pursue. Maps were horizontally flipped for half of the participants to control for any left-right biases.

We then applied the same computational modeling framework to our human participant data, which allowed us to test which representational models and reliance parameters best fit the data.

##### 2.4. Results

Human participants preferred to choose the reward that was hidden from their competitor on the Center-Wall (198/200 participants, $p_{binom} < 0.001$), Open-Hidden (199/200 participants, $p_{binom} < 0.001$), and Transparent-Hidden Routes paradigms (190/200 participants, $p_{binom} < 0.001$). On the Hidden-Hidden paradigm, participants chose at chance (109/200 participants, $p_{binom} = 0.23$). On the Open-Transparent paradigm, participants displayed a preference to choose the reward behind the transparent barrier over the reward out in the open (147/200 participants, $p_{binom} < 0.001$). Although (according to the instructions) the competitor could see both rewards and would therefore steal the reward from the participant, participants' preference for choosing the reward behind the transparent barrier could reflect a desire to have a physical barrier, even a transparent one, rather than nothing between them and the competitor. Human participants might heavily weigh the physical cost to the competitor of navigating around the transparent barrier to reach the candy. Overall, these results confirm that, on these tasks, adult humans use their understanding of how an agent's visual perspective affects their knowledge to choose the rewards that were hidden from their competitor.

##### 2.5. Discussion

Adult human behavior was best described by the *Ignorance rep.* model, but this model, like all models, failed to predict adult humans' preference on the Open-Transparent paradigm for the food behind the transparent barrier. This paradigm is responsible for most of the gap ($\Delta LL = -14.1$ out of the total $\Delta LL = -15.2$) between the ceiling of a model's maximum possible explanatory power ($LL = -10.0$) and the *Ignorance rep.* model's explanatory power (LL=$-25.2$).

#### 3. Additional results

##### 3.1. Walk-through of qualitative model behavior

Here, we give a qualitative description of how each model behaves (assuming full reliance, $r = 1$).

As shown in Fig. 3 in the main text, the egocentric model (*Physical planner*) and the behavioral *Food-directed behavior* model predicted chance performance on all paradigms. On the Open-Transparent paradigm, the behavioral (*Location-directed behavior* model predicted that the competitor would prefer to pursue the food out in the open to avoid the cost of navigating around the transparent barrier, and therefore predicted that the subject would slightly prefer the food behind the transparent barrier. With $r = 1$, this preference was 0.57.

The mentalistic models revealed mixed performance. Like the egocentric and behavioral models, the *True belief* model failed to produce the directional preferences displayed by NHPs on the Center-Wall, Open-Hidden, and Transparent-Hidden Routes paradigms. However, on the Center-Wall paradigm, the *True belief* model predicts that the competitor is slightly more likely ($p = 0.53$) to pursue the food in the center than behind the wall because the food at the center is marginally closer to the competitor. This is what results in the 0.53 preference in the subject for the food behind the barrier. On the Open-Transparent paradigm, like the *Location-directed behavior* model, the

| | *Physical planner* | *Food-directed* | *Location-directed* | *True belief* | *Partial* | *Uncertainty* | *Ignorance* | *NHPs* |
|---|---|---|---|---|---|---|---|---|
| CW | 0.51 | 0.50 | 0.51 | 0.53 | 1.0 | 0.76 | 1.0 | 0.83 |
| OH | 0.49 | 0.50 | 0.50 | 0.51 | 1.0 | 0.75 | 1.0 | 0.67 |
| THR | 0.51 | 0.50 | 0.50 | 0.51 | 0.83 | 0.85 | 0.91 | 0.57 |
| OT | 0.49 | 0.50 | 0.57 | 0.57 | 0.57 | 0.57 | 0.57 | 0.55 |
| HH | 0.50 | 0.50 | 0.50 | 0.49 | 0.50 | 0.50 | 0.50 | – |

Table S1: Table of the results presented in Fig. 4 in main. Model behavior under full reliance ($r = 1$). For the three paradigms with hidden food items, the probabilities shown are of choosing the hidden food. In the two control paradigms (Open-Transparent and Hidden-Hidden), the probabilities represent the chance of going for the right-most food shown in Fig. 2 in main. CW: Center-Wall. OH: Open-Hidden. THR: Transparent-Hidden Routes. OT: Open Transparent. HH: Hidden-Hidden. The exact effect size on the Hidden-Hidden paradigm in Hare et al. 2000 could not be recovered (see 4.1).

*True belief* predicts that the subject would go for the food behind the transparent barrier with probability 0.57. All mentalistic models make this prediction on Open-Transparent.

In contrast, the *Uncertainty rep.* model predicts moderate effect sizes in the directional paradigms (76%, 75%, and 85% for Center-Wall, Open-Hidden, and Transparent-Hidden Routes, respectively). Because this model integrates over two epistemic hypotheses (that the competitor does or doesn't know about the hidden food), it essentially averages over a 100% and 50% chance (produced from the MDPs representing the competitor) that the competitor will pursue the hidden food in the Center-Wall and Open-Hidden paradigms, producing the prediction of 75%. In the Transparent-Hidden Paradigm, the probability of detection along the hidden route follows $p \sim Beta(\alpha = 9, \beta = 1)/2$ and $p \sim Beta(\alpha = 9, \beta = 1)$ along the transparent route (see SM Section 1.1). 85% of the time, the subject will choose the hidden route.

Finally, the *Partial rep.* and *Ignorance rep.* models predict near-ceiling performance in the Center-Wall and Open-Hidden paradigms. On the Transparent-Hidden Routes paradigm, the *Ignorance rep.* and *Partial rep.* models predicted effect sizes of 91% and 83%, respectively. The *Partial rep.* model predicts a smaller effect size because it cannot predict anything about the competitor's probability of pursuing the food on the hidden side (since its representation of the competitor's mind is missing the hidden side of the map). Therefore, the *Partial rep.* cannot predict that the competitor will not go there, and its preference for the hidden side is weaker than the *Ignorance rep.* model's preference.

*3.2. Walk-through of models and comparison to human and NHP empirical data*

Here, we describe how the models behave when assuming full reliance ($r = 1$) and compare them first against human and then NHP empirical data. Fig. 4 in the main text shows each model's performance on each paradigm overlaid with human and NHP observed results.

Qualitatively, the more complex models better match human behavior than the simpler models. The full *Ignorance rep.* model best matches human performance, capturing human performance particularly well on the Transparent-Hidden Routes paradigm, which no other model captures as well. *Partial rep.* makes very similar predictions to the *Ignorance rep.* model, but predicts slightly weaker effects on Transparent-Hidden Routes and therefore underestimates human performance on this paradigm. (On Transparent-Hidden Routes, humans choose the hidden route 95% of the time. The *Ignorance rep.* model similarly chooses the hidden route model 91% of the time, while the *Partial rep.* model chooses it only 83% of the time.) The *Uncertain rep.* model predicts effect sizes that are too small on Center-Wall, Open-Hidden, and Transparent-Hidden Routes paradigms (76%, 75%, and 85%, respectively). The *True belief*, *Location-directed*, *Food-directed*, and *Physical planner* models all failed to match human preferences on these directional paradigms. The result that the *Ignorance rep.* model best describes the behavior of adult humans helps validate that the models and human participants are behaving as expected on these tasks.

Notably, on the Open-Transparent paradigm, human participants favored the food behind the transparent barrier. This suggests that humans might weigh the physical cost to the competitor more strongly, preferring to have a physical barrier between them and the source of danger, regardless of whether that barrier is transparent or not. Because of the physical cost imposed on the competitor for navigating around the transparent barrier, the *Ignorance rep.*, *Uncertainty rep.*, *Partial rep.*, *True belief*, and *Location-directed* models predicted very slight preferences for the food behind the transparent barrier. Increasing the movement costs to the competitor or the sensitivity to those costs (via the $\tau_{choice}$ parameter) would enable these models to better capture human behavior on this paradigm.

Turning from human to NHP behavior, the three most complex models (*Ignorance rep.*, *Uncertainty rep.*, and *Partial rep.*) were all able to capture the direction of NHP behavior (Fig. 3 in main), but not the exact effect sizes.

These complex models generally tended to produce stronger effect sizes on the directional paradigms (Center-Wall, Open-Hidden, and Transparent-Hidden Routes) than NHPs displayed. The only exception is that the *Uncertainty rep.* model predicted a smaller effect size than NHPs showed on the Center-Wall paradigm: *Uncertainty rep.* choose the food in the center 76% of the time, while NHPs only chose it 83% of the time. On every other directional paradigm, the *Uncertainty rep.*, *Partial rep.*, and *Ignorance rep.* models predicted effect sizes that were too large. Finally, the simplest model in the mentalistic family (*True belief*) and the behavioral (*Location-directed Food-directed*), and the ego-centric *Physical planner* models all failed to predict NHP preferences on the three directional paradigms. However, all seven models were able to capture NHPs behavior on the two paradigms where NHPs performed at chance.

###### *3.2.1. RMSE analysis*

To capture these observations quantitatively, we calculated the squared error of each model against NHP and human data on each paradigm, and then averaged across paradigms to yield MSE. The Hidden-Hidden control paradigm is the only one for which no choice data are available (see SM 4.1). Conveniently, all models make the same prediction for this paradigm: that the agent will choose at chance (50-50) between the two foods. For the purposes of the visualization in Fig. 4 in main and for the MSE calculation, we assumed that the NHPs also choose at chance. Since all models, then, have zero error on this paradigm, this assumption does not favor any model over the others. MSEs are reported below.

| **Model** | **MSE NHPs** | **MSE Humans** |
|---|---|---|
| Physical planner | 0.17 | 0.39 |
| Food-directed | 0.17 | 0.39 |
| Location-directed | 0.17 | 0.38 |
| True belief | 0.16 | 0.37 |
| Partial rep. | 0.20 | 0.092 |
| Uncertainty rep. | 0.13 | 0.17 |
| Ignorance rep. | 0.23 | 0.077 |

Table S2: MSEs between models (assuming full reliance) and empirical data.

##### *3.3. Explanatory power as a function of reliance*

The results in the previous section depended on assuming that NHPs and humans always rely on the model's posited representation. Here, we examine the effects of varying the degree of reliance on each model. Figure S2 shows each model's explanatory power as a function of reliance.

The *Physical planner* model is insensitive to the reliance parameter because, by design, it does not have any representation of the competitor. The behavioral *Food-directed behavior* model performed very similarly to the *Physical planner* model, and therefore, reliance affected explanatory power minimally, resulting in a nearly flat line in Figure S2. The behavioral *Location-directed behavior* and the mentalistic *True belief* model showed a similar structure to each other: the stronger the assumption that primates always use these representations (the higher the value of $r$), the better these models could explain the empirical data. The other two models in the mentalistic family were best able to explain the empirical data under the assumption that NHPs rely on their social representations most, but critically, not all the time: $r = 0.58$ for *Uncertainty rep.* and $r = 0.53$ for *Partial rep.*. Finally, the *Ignorance rep.* model's ability to explain the empirical data peaked when positing that NHPs use their full theory of mind about a third of the time ($r = 0.35$), and rapidly decreased when it was posited that subjects always relied on their underlying social representations, because the model predicted larger effect sizes than the empirical data showed. By way of validation, the *Ignorance rep.* model's ability to explain human data reached its peak at $r = 0.99$, because the human empirical data showed the large effect that the *Ignorance rep.* model predicted. (At reliance $r = 1.0$, the *Ignorance rep.* predicted perfect performance and thus could not tolerate any noise.) The maximum of each curve in Fig. S2 is plotted in Fig. 5 in the main text.

##### *3.4. Model behavior under optimal reliance*

Table S3 depicts the behavior of each model under optimal reliance for maximizing its explanatory power over the NHP data. For convenience, NHP behavioral data is also displayed.

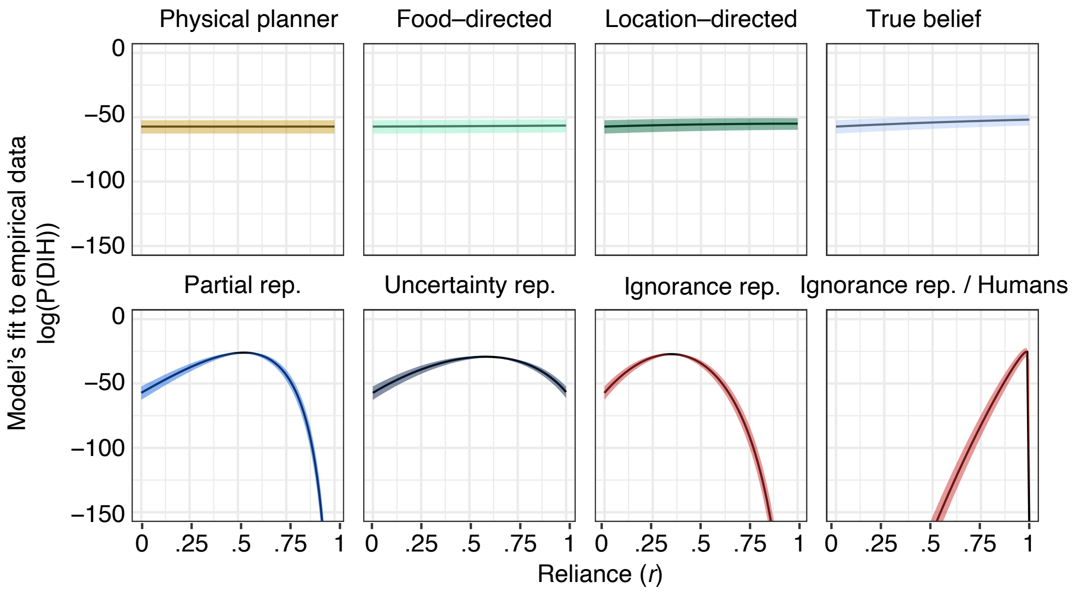

Figure S2: Model explanatory power as a function of the hypothesized reliance $r$. The x-axis shows reliance. When $r = 0$, the subjects never rely on the posited social representations and when $r = 1$, the subjects always rely on the model's posited social representations. The y-axis shows the probability (in log-space) of the model replicating the experimental data under that reliance value. All models produce the same likelihood (for the NHP data, LL=−57.5) at $r = 0$ because at zero reliance, they all simply express an egocentric planner.

| | *Physical planner* | *Food-directed* | *Location-directed* | *True belief* | *Partial* | *Uncertainty* | *Ignorance* | *NHPs* |
|---|---|---|---|---|---|---|---|---|
| CW | 0.51 | 0.50 | 0.51 | 0.53 | 0.69 | 0.65 | 0.68 | 0.83 |
| OH | 0.49 | 0.50 | 0.50 | 0.51 | 0.68 | 0.64 | 0.67 | 0.67 |
| THR | 0.51 | 0.50 | 0.50 | 0.51 | 0.64 | 0.67 | 0.66 | 0.57 |
| OT | 0.49 | 0.50 | 0.57 | 0.57 | 0.53 | 0.54 | 0.52 | 0.55 |
| HH | 0.50 | 0.50 | 0.50 | 0.49 | 0.50 | 0.50 | 0.50 | – |

Table S3: Model behavior under best reliance. For the three paradigms with hidden food items, the probabilities shown are of choosing the hidden food. In the two control paradigms (Open-Transparent and Hidden-Hidden), the probabilities represent the chance of going for the right-most food shown in Fig. 2 in main. CW: Center-Wall. OH: Open-Hidden. THR: Transparent-Hidden Routes. OT: Open Transparent. HH: Hidden-Hidden. The exact effect size on the Hidden-Hidden paradigm in Hare et al. 2000 could not be recovered (see 4.1).

#### 4. Additional methods

##### *4.1. Behavioral data*

Our models aim to capture the effect sizes displayed by non-human primate behavior reported in the experiments in Table 1 in Main. For Bräuer et al. (2007), Hare et al. (2006), and the Hidden-Hidden conditions of Hare et al. (2000) E3 and E4, data on the proportions of choices are incomplete or unavailable, and thus we are unable to include these experiments in our quantitative model evaluations. For most other studies, the proportions of choices are clearly reported. The experiments for which we could most easily recover the choice data from the reported proportions and number of trials were Hare et al. (2000) Experiments 1 (45 choices of the hidden food out of 54 trials), E2 (20 out of 27), E3 Open-Hidden condition (52 out of 83), E4 Open-Hidden condition (62 out of 108), and Canteloup et al. (2016) Experiment 1, Condition 2 (95 out of 125). Choice data for Hare et al. (2000) E5 and Melis et al. (2006) E1 were also recoverable, but required making a few reasonable assumptions. In Hare et al. (2000) E5, we assumed a total of 11 subjects with 12 trials per subject based on indirect details elsewhere in the paper, enabling the recovery of choice data (73 out of 132). For Melis et al. (2006) E1, by assuming that trials in which a subject did not make a choice were excluded and by reading mean choices from Figure 2, we recovered the choice data (72 out of 126). When two versions of the same experiment were available that differed only in the timing of the competitor's release (Canteloup et al. 2016, E1; Hare et al. 2000, E5), we used the version with a longer delay, which better controls for the possibility that the subject's choice could be influenced by first observing the competitor's direction of movement. Table S4 shows a summary of the recovered choice data.

| Paradigms | Paper | Exp | Result |
|---|---|---|---|
| Center-Wall | Hare et al., 2000 | E1 | 45/54 |
| Open-Hidden | Hare et al., 2000 | E2 | 20/27 |
| Open-Hidden | Hare et al., 2000 | E3 | 52/83 |
| Open-Hidden | Hare et al., 2000 | E4 | 62/108 |
| Open-Hidden | Canteloup et al., 2016 | E1-C2 | 95/125 |
| Open-Transparent | Hare et al., 2000 | E5 | 73/132 |
| Transparent-Hidden Routes | Melis et al., 2006 | E1 | 72/126 |

Table S4: NHP data used in quantitative model evaluation.

*4.2. Coding qualitative model predictions*

Because each computational model predicts a continuous effect size (rather than an absolute success or failure), we coded model behavior as predicting chance performance when its preference for both food rewards was below a threshold $t$, and as making a directional prediction otherwise. In the relevant literature, effects are considered to be directional when they significantly differ from chance (0.5). How much deviation from 0.5 is significant depends on the sample size.

The experiments considered had sample sizes given in the denominator of the results column in Table S4. Assuming a two-tailed binomial test against chance ($p = 0.5$) at significance level $\alpha = 0.05$, the average effect size detectable by these experiments was 0.61. Therefore, we set the threshold value $t = 0.6$.
